# Identification of molecular determinants governing KIF4 cargo recognition of NTCP and their role in HBV/HDV infection

**DOI:** 10.64898/2026.02.26.708216

**Authors:** Sameh A. Gad, Doaa A. Abo Elwafa, Masaaki Toyama, Mitsunori Ikeguchi, Ikuo Shoji, Kazuaki Chayama, Masanori Isogawa, Takaji Wakita, Hussein H. Aly

## Abstract

The surface availability of the sodium taurocholate co-transporting polypeptide (NTCP), which serves as entry receptor for Hepatitis B virus (HBV) and Hepatitis D virus (HDV), is a key determinant of HBV/HDV infectivity. We previously identified the Kinesin motor protein KIF4 as a key regulator of NTCP trafficking to the plasma membrane. However, the molecular mechanism by which NTCP is recognized as a KIF4 transport cargo has remained undefined. Here, we identified a leucine-rich motif, LLLI (aa 298–301), within the C-terminal transmembrane domain of NTCP as an essential determinant for KIF4 transport. Co-immunoprecipitation analyses revealed its importance for NTCP interaction with KIF4. This motif is conserved in another established KIF4 cargo, integrin β1 (ITGB1), and mutation of the corresponding residues similarly disrupted ITGB1–KIF4 interaction, indicating a shared cargo recognition mechanism employed by KIF4. Functionally, disruption of the NTCP leucine-rich motif markedly impaired NTCP surface localization without affecting total protein expression, resulting in severe attenuation of HBV and HDV infection. Structural modeling and computational protein–protein interaction analysis indicated that the mutation does not alter NTCP structure or NTCP/preS1 binding affinity, supporting a trafficking-specific defect. Substitution mutations identified L298 and L299 as the minimal residues required for interaction with KIF4. Our findings define a recognition motif that regulates KIF4-dependent cargo identification and surface trafficking. It also underscores the potential of targeting KIF4-mediated NTCP recognition and transport to inhibit HBV and HDV entry.

**Importance:** Hepatitis B and D viruses require the hepatocyte surface receptor NTCP for entry. We identified a conserved leucine-rich motif (LLLI, aa 298–301) in NTCP that is necessary for its recognition by the kinesin motor KIF4; disrupting this motif selectively impairs NTCP surface trafficking and markedly attenuates HBV/HDV infection without reducing total NTCP or intrinsic receptor function. Importantly, we identified a similar motif in integrin β1 (ITGB1; aa 747–750) that is required for interaction with KIF4, revealing a similar cargo selection mechanism for different proteins. Scientifically, these findings define a motif-derived cargo-selection determinant implied by KIF4, advancing a general principle for kinesin-dependent cargo recognition. More broadly, this work highlights a possible host-directed antiviral approach that targets cargo-selection interfaces and provides a conceptual framework for manipulating kinesin-mediated trafficking of clinically relevant membrane proteins beyond NTCP, with implications for receptor regulation and membrane protein therapeutics.

## Introduction

Chronic hepatitis B virus (HBV) infection remains a major global health burden, affecting nearly 1.5 million new infections/year with 254 million individuals suffering from chronic HBV worldwide. Chronic HBV patients may progress to complications from the progressive liver disease such as cirrhosis, and hepatocellular carcinoma (1). Clinical outcomes of HBV patients worsen substantially in the presence of hepatitis D virus (HDV), a defective satellite virus that requires HBV envelope proteins for propagation. HDV coinfection accelerates liver inflammation, increases the risk of fulminant hepatitis, and markedly elevates the likelihood of end-stage liver disease (2). A pivotal step in the life cycle of both HBV and HDV is their shared dependence on the sodium taurocholate co-transporting polypeptide (NTCP), a bile acid transporter that functions as the essential entry receptor on hepatocytes (3). Understanding how NTCP is trafficked and presented on the cell surface is therefore central to elucidating the mechanisms governing HBV/HDV infection (4).

Kinesin family member 4 (KIF4) is a microtubule-based motor protein involved in intracellular transport, chromosome segregation, and cytoskeletal organization (5). Beyond its canonical roles in mitosis and nuclear dynamics, emerging evidence indicates that KIF4 also functions in cytoplasmic vesicular trafficking and regulates the transport and surface localization of specific membrane proteins (6–8). Our previous findings revealed that KIF4 interacts with NTCP and facilitates its anterograde transport to the plasma membrane, thereby enabling the receptor’s availability for HBV/HDV attachment and viral entry (4). These observations positioned KIF4 as a previously unrecognized host factor that modulates viral susceptibility by controlling NTCP surface expression (4, 9).

Although kinesins share highly conserved motor domains, cargo specificity is primarily determined by divergent C terminal tail domain that engage adaptor proteins, recognize discrete cargo binding motifs, and integrate regulatory inputs such as post translational modifications and scaffolding complex assembly (10–13). For many kinesins, such as KIF1A, KIF5, and KIF3 family members, structural studies have identified short linear motifs or coiled-coil regions that mediate selective cargo recognition (11, 12, 14, 15). However, the molecular basis by which KIF4 identifies and transports its physiological cargoes remains poorly defined. Although, we clearly demonstrated the interaction between KIF4 and NTCP by pull-down assays and intracellular localization studies, recombinant KIF4 and NTCP failed to exhibit a direct physical association in vitro. This observation suggests that the KIF4–NTCP interaction is likely facilitated by specific adaptor proteins that mediate or regulate this association within the cellular environment (4). Only limited information exists regarding KIF4’s tail-domain interactome, and no detailed determinants have been identified for its cargo selection of membrane proteins such as NTCP. This gap in knowledge reduces our ability to understand how KIF4-dependent cargo trafficking is regulated and how it may be selectively modulated.

Defining the molecular determinants that govern KIF4–cargo recognition, particularly in the context of NTCP, holds substantial therapeutic relevance. Because NTCP surface availability is a rate-limiting step for HBV/HDV entry (3, 4), elucidating how KIF4 recognizes and transports NTCP could reveal new intervention points for antiviral drug development (4, 9). Targeting the specific interaction interface—rather than globally inhibiting kinesin activity—offers the possibility of selectively disrupting viral entry while preserving essential cellular transport processes. A mechanistic understanding of this cargo recognition pathway may therefore enable the rational design of small molecules or peptides that block NTCP trafficking, providing a novel host-directed strategy to prevent HBV/HDV infection.

In this study, we identify the molecular basis by which the kinesin motor protein KIF4 recognizes (NTCP and ITGB1) as a transport cargo and regulates its delivery to the hepatocyte plasma membrane. We show that KIF4-mediated transport requires the leucine-rich motif (LLLI) present at the C-terminal transmembrane domain of both membrane proteins. Disruption of this motif abolishes the interaction between KIF4 and its cargo (NTCP or ITGB1), markedly reduces the surface expression of both proteins without affecting total protein levels. As a result, suppression of NTCP surface expression severely impaired HBV and HDV entry. Together, these findings define a previously unrecognized kinesin cargo-recognition motif that controls NTCP surface localization and represents a critical regulatory node in HBV and HDV infection.

## Materials and Methods

### Cell lines and peptides

HepG2 were cultured with DMEM/F-12 + GlutaMax medium (Gibco) supplemented with 10% fetal bovine serum (FBS), 10 mM HEPES, 100 unit/ml penicillin, 100 μg/ml streptomycin, and 5 μg/ml insulin (4). HEK293FT cells were cultured in DMEM, high glucose supplemented with 10% FBS, sodium pyruvate (1 mM), MEM non-essential amino acids (1X), penicillin 100 unit/ml penicillin, 100 μg/ml streptomycin. All cell lines used in this work were incubated at 37 °C in the presence of 5% CO_2_ (9). Fluorescent HBV preS1 probe, a peptide spanning the 2-48 amino acids of the L-HBsAg preS1 region with myristoylation and 6-carboxytetramethylrhodamine (TAMRA) labeling at N- and C-termini, respectively, was synthesized by Scrum, Inc.

### Plasmid Construction

The N-terminal 6×Myc-tagged KIF4 expression construct in the pIRESpuro3 vector was kindly provided by Dr. Toru Hirota (Cancer Institute of the Japanese Foundation for Cancer Research, JFCR) (16). C-terminally HA-tagged full-length NTCP (GenBank Accession No. NM_003049.4), cloned into the CSII-EF-MCS vector at the *XhoI* (5′) and *XbaI* (3′) sites and provided by Dr. Hiroyuki Miyoshi (RIKEN BioResource Research Center, Japan), was subcloned into the pcDNA3.1 expression vector. The NTCP-HA open reading frame (ORF) was excised by *XhoI* and *XbaI* and ligated into *XhoI/XbaI*-digested pcDNA3.1 to generate full-length NTCP-HA in pcDNA3.1.

### Generation of full-length NTCP-HA alanine-substitution mutants

The full-length NTCP-HA construct carrying four alanine substitutions (L298A, L299A, L300A, and I301A) was generated by introducing these mutations into wild-type (WT) NTCP-HA using a two-step PCR approach. Two overlapping PCR fragments were first amplified from WT NTCP-HA using the following primer pairs:

- **Fragment 1:** Forward 5′-TAACGCTCGAGATGGAGGCCCACAAC-3′ Reverse 5′- CTCATAGCACCAAAATATGGCGGCGGCGGCGGCCCCTTCTCCAAGCTGGAAA ATC-3′
- **Fragment 2:** Forward 5′- GATTTTCCAGCTTGGAGAAGGGGCCGCCGCCGCCGCCATATTTTGGTGCTATGAG-3′ Reverse 5′-GGCCCTCTAGACTAAGCGTAATCTGGAAC-3′

A third PCR reaction was then performed using the two overlapping amplicons as templates, with the forward primer of Fragment 1 and the reverse primer of Fragment 2. The resulting full-length mutated NTCP-HA fragment was digested with *XhoI* and *XbaI* and ligated into *XhoI/XbaI*-digested pcDNA3.1.

### Generation of NTCP truncated constructs

Two N-terminal NTCP truncations, NTCP-HA (aa 1–233) and NTCP-HA (aa 1–117), were generated by PCR amplification of WT NTCP-HA using:

- **Forward primer** (both constructs): 5′-TGCCAGGTACCATGGAGGCCCACAAC-3′
- **Reverse primers:** NTCP-HA (aa 1–233): 5′-TCGGCTCTAGACTAAGCGTAATCTGGAACATCGTATGGGTAGCCAATAAAAGGCATCAGGG-3′ NTCP-HA (aa 1–117): 5′-TCGGCTCTAGACTAAGCGTAATCTGGAACATCGTATGGGTAGTTCATGTCCCCCTTCATGG-3′

PCR products were digested with *KpnI* and *XbaI* and subcloned into *KpnI/XbaI*-digested pcDNA3.1.

A C-terminal region construct NTCP-HA (aa 234–349) was similarly generated using:

- Forward 5′-TGCCAGGTACCATGTTTCTGCTGGGTTATG-3′
- Reverse 5′-TCGGCTCTAGACTAAGCGTAATCTGGAACATCGTATGGGTAGGCTGTGCAAGGGGAGCAGT-3′ and inserted into *KpnI/XbaI*-digested pcDNA3.1.

### Generation of NTCP-HA (aa 234–349) constructs carrying alanine-substitution mutations

The truncated NTCP-HA (aa 234–349) construct carrying the quadruple mutation (L298A/L299A/L300A/I301A) was generated using the same overlapping PCR strategy employed for full-length NTCP. Two PCR fragments were amplified from WT NTCP-HA (aa 234–349) using the primer pairs:

- **Fragment 1:** Forward 5′-TGCCAGGTACCATGTTTCTGCTGGGTTATG-3′ Reverse 5′-CTCATAGCACCAAAATATGGCGGCGGCGGCGGCCCCTTCTCCAAGCTGGAAA ATC-3′
- **Fragment 2:** Forward 5′-GATTTTCCAGCTTGGAGAAGGGGCCGCCGCCGCCGCCATATTTTGGTGCTATGAG-3′ Reverse 5′-GGCCCTCTAGACTAAGCGTAATCTGGAAC-3′

The overlap-extension PCR product was digested with *KpnI* and *XbaI* and subcloned into *KpnI/XbaI*-digested pcDNA3.1.

### Generation of NTCP-HA (aa 234–349) single alanine-substitution mutants

Single alanine-substitution mutants of NTCP-HA (aa 234–349) (L298A, L299A, L300A, or I301A) were generated using the same overlap-extension PCR strategy, with WT NTCP-HA (aa 234–349) as the template.

For each mutant, two PCR reactions were performed as follows:

- **L298A mutant:** PCR reaction 1: Forward: 5′-TGCCAGGTACCATGTTTCTGCTGGGTTATG-3′ Reverse: 5′-CTCATAGCACCAAAATATGGCAATGAGGAGGGCCCCTTCTCCAAGCTGGAAA ATC-3′ PCR reaction 2: Forward: 5′-GATTTTCCAGCTTGGAGAAGGGGCCCTCCTCATTGCCATATTTTGGTGCTATGAG-3′ Reverse: 5′-GGCCCTCTAGACTAAGCGTAATCTGGAAC-3′
- **L299A mutant:** PCR reaction 1: Forward: 5′-TGCCAGGTACCATGTTTCTGCTGGGTTATG-3′ Reverse: 5′-CTCATAGCACCAAAATATGGCAATGAGGGCAAGCCCTTCTCCAAGCTGGAAA ATC-3′ PCR reaction 2: Forward: 5′-GATTTTCCAGCTTGGAGAAGGGCTTGCCCTCATTGCCATATTTTGGTGCTATGAG-3′ Reverse: 5′-GGCCCTCTAGACTAAGCGTAATCTGGAAC-3′
- **L300A mutant:** PCR reaction 1: Forward: 5′-TGCCAGGTACCATGTTTCTGCTGGGTTATG-3′ Reverse: 5′-CTCATAGCACCAAAATATGGCAATGGCGAGAAGCCCTTCTCCAAGCTGGAAA ATC-3′ PCR reaction 2: Forward: 5′-GATTTTCCAGCTTGGAGAAGGGCTTCTCGCCATTGCCATATTTTGGTGCTATGAG-3′ Reverse: 5′-GGCCCTCTAGACTAAGCGTAATCTGGAAC-3′
- **I301A mutant:** PCR reaction 1: Forward: 5′-TGCCAGGTACCATGTTTCTGCTGGGTTATG-3′ Reverse: 5′-CTCATAGCACCAAAATATGGCGGCGAGGAGAAGCCCTTCTCCAAGCTGGAAA ATC-3′ PCR reaction 2: Forward: 5′-GATTTTCCAGCTTGGAGAAGGGCTTCTCCTCGCCGCCATATTTTGGTGCTATGAG-3′ Reverse: 5′-GGCCCTCTAGACTAAGCGTAATCTGGAAC-3′

For each mutant, a third PCR reaction was performed using the products of the two reactions as templates, with the forward primer from PCR reaction 1 and the reverse primer from PCR reaction 2. The final PCR products were digested with *KpnI* and *XbaI* and ligated into *KpnI*/*XbaI*-digested pcDNA3.1.

### Plasmid transfection

Plasmid transfections in HEK293FT cells were performed using Lipofectamine 2000, whereas transfections in HepG2 cells were carried out using Lipofectamine 3000, in accordance with the manufacturer’s instructions.

### Immunoblot analysis

Protein expressions were analyzed by immunoblotting as previously described (17). The primary antibodies used were mouse monoclonal anti-Myc (Santa Cruz Biotechnology, sc-40), anti–E-cadherin (Santa Cruz Biotechnology, sc-8426), and anti–β-actin (Sigma-Aldrich, A5441), as well as rabbit polyclonal anti-HA (Sigma-Aldrich, H6908). To assess NTCP expression, protein samples were treated with Peptide-N-Glycosidase F (PNGase F; 250 U) prior to SDS–PAGE to remove N-linked glycans from glycosylated NTCP, as previously reported (9).

### Isolation of the cell surface proteins

Cell surface proteins were isolated by surface biotinylation followed by streptavidin–agarose affinity purification, as previously described (4). The recovered cell surface fractions were subsequently analyzed by immunoblotting as described above. E-cadherin (CDH1) was used as an internal loading control to verify the enrichment and integrity of the surface protein fraction (18).

### Co-immunoprecipitation (Co-IP) assay

HEK293FT cells were co-transfected with the indicated HA-tagged NTCP or HA-tagged ITGB1 expression plasmids together with a Myc-tagged KIF4 expression plasmid at a 1:1 DNA ratio. At 72 h post-transfection, cells were harvested and lysed, and protein complexes were subjected to co-immunoprecipitation (co-IP) using a mouse monoclonal anti-Myc antibody (Santa Cruz Biotechnology, sc-40). Mouse normal IgG was used in parallel as a negative control to assess assay specificity. Following immunoprecipitation, the recovered complexes were analyzed by immunoblotting to detect co-immunoprecipitated HA-tagged NTCP or HA-tagged ITGB1 in the pull-down fraction. Cell lysis and co-IP procedures were performed using the Pierce Co-Immunoprecipitation Kit (Thermo Fisher Scientific, 26149) in accordance with the manufacturer’s instructions.

### HBV preS1 peptide binding assay

Cell surface attachment of the N-terminally myristoylated and C-terminally TAMRA-labeled HBV preS1 peptide was assessed in HepG2 cells transiently expressing hNTCP as previously described (9).

### HBV infection assay

HBV viral inoculum was prepared from the culture supernatant of HepAD38.7-Tet cells as previously described (19). HepG2 cells transiently expressing hNTCP were inoculated with HBV at a multiplicity corresponding to 10,000 genome equivalents (GEq) per cell in the presence of 4% PEG8000. Following a 16-h incubation, unbound viral particles were removed by extensive washing, and cells were maintained in culture for an additional 7 days. HBV infection was assessed by quantifying secreted HBeAg using ELISA and intracellular HBV pgRNA levels by real-time RT-PCR.

### HDV infection assay

HDV used in infection experiments was generated from the culture supernatant of Huh7 cells co-transfected with pSVLD3 and pT7HB2.7 plasmids, as previously described (20, 21). HepG2 cells transiently expressing hNTCP were inoculated with HDV at 100 GEq per cell in the presence of 5% PEG8000. After a 16-h incubation, unbound virus was removed by extensive washing, and the cells were maintained in culture for an additional 6 days. HDV infection was evaluated by detection of HDAg using immunofluorescence analysis.

### Immunofluorescence (IF) assay

Immunofluorescence staining was performed as previously described (4). Briefly, cells were fixed with 4% paraformaldehyde and permeabilized with 0.3% Triton X-100. The samples were then incubated with a rabbit anti-HDAg primary antibody (developed by Scrum), followed by incubation with an Alexa Fluor 555–conjugated secondary antibody. Nuclei were counterstained with DAPI. Fluorescence signals were examined and imaged using a fluorescence microscope.

### Structural prediction and comparative analysis of NTCP

The three-dimensional structures of wild-type NTCP and the NTCP mutants were predicted using AlphaFold 3.0.0 installed on a local server. The default databases were used, including the Google DeepMind–curated versions of BFD, MGnify (v2022_05), UniProt (2021_04), UniRef90 (2022_05), PDB, and PDB_seqres (2022_09_08). The wild-type NTCP sequence was obtained from UniProt (Q14973). For the mutants L298A, L299A, L300A, I301A, and the quadruple mutant L298A/L299A/L300A/I301A, the corresponding residues in the sequence were modified accordingly. For each prediction, a single random seed was used, and among the five generated output models, the structure with the highest confidence score was selected. RMSD calculations were performed using the Cα atoms of residues 22–309.

### Amino acid sequence alignment

Multiple sequence alignments were performed for the C-terminal 116 amino acids of NTCP (aa 234–349) in comparison with human ITGB1, as well as for the NTCP C-terminal transmembrane sequence (aa 294–305) across HBV-susceptible species. Alignments were generated using the Clustal Omega algorithm (EMBL-EBI web server) with default parameters. Graphical representations of sequence conservation and alignment annotations were prepared using ESPript 3.2 (22–24).

### Statistical analysis

All experiments were performed in triplicate, and data represent the mean ± standard deviation (SD) from three independent experiments. Statistical significance was assessed using an unpaired two-tailed Student’s *t* test (**, P < 0.01; ***, P < 0.001; NS, not significant).

## Results

### The C-terminal Domain of NTCP is crucial for interaction with KIF4

We previously demonstrated that KIF4 mediates NTCP trafficking to the hepatocyte cell surface(4). To further clarify the mechanism by which KIF4 recognizes and interacts with its cargo, we constructed expression plasmids encoding C-terminal HA-tagged full-length NTCP, as well as NTCP truncation mutants containing either the first 117 or 233 amino acids (Fig. 1A). Each NTCP construct was separately co-transfected with a Myc-tagged KIF4 expression plasmid into HEK293FT cells and incubated for 72 hours, after which co-immunoprecipitation (co-IP) assays were performed. Immunoprecipitation with the anti-Myc antibody resulted in efficient recovery of KIF4, while no KIF4 was detected in samples processed with the isotype control antibody, thereby validating the specificity of the assay (Fig. 1B). Notably, co-immunoprecipitation analysis demonstrated that KIF4 selectively associates with the full-length NTCP, whereas truncated NTCP variants failed to associate with KIF4 (Fig. 1B), indicating that the interaction between NTCP and KIF4 requires the C-terminal region of NTCP.

**FIG 1.**
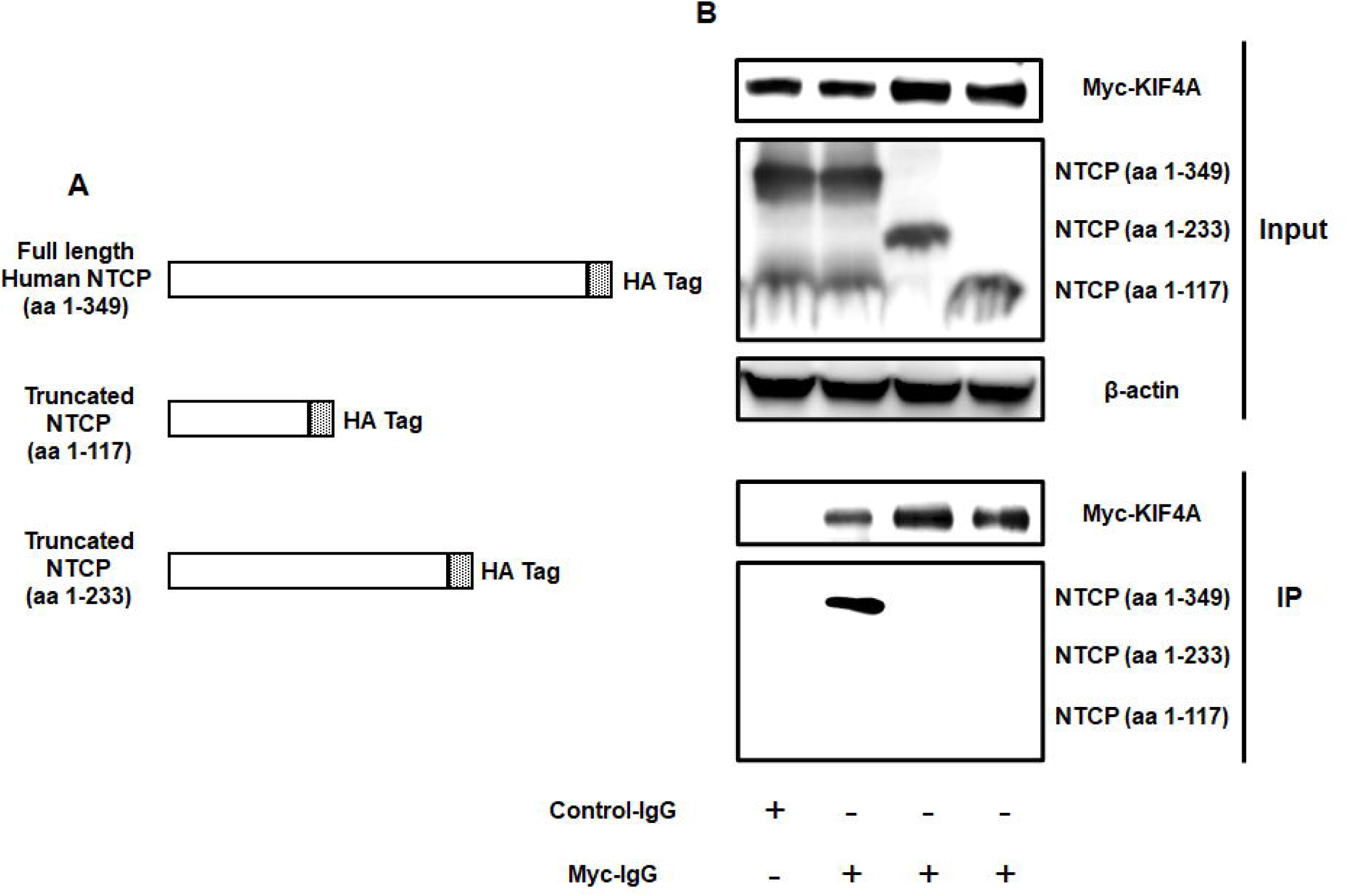
KIF4 interacts with the C-terminal region of NTCP. **(A)** Schematic representation of C-terminally HA-tagged full-length NTCP (aa 1–349) and C-terminal truncation mutants comprising the N-terminal 117 (aa 1–117) or 233 (aa 1–233) amino acids. **(B)** HEK293FT cells were co-transfected with expression plasmids encoding Myc-tagged KIF4 and C-terminally HA-tagged full-length NTCP or the indicated NTCP truncation mutants at a 1:1 ratio. At 72 h post-transfection, cells were lysed, and 10% of each lysate was analyzed by immunoblotting to detect Myc-KIF4 (*Input; uppermost panel*), HA-NTCP (*Input; middle panel*), and β-actin (loading control) (*Input; lowermost panel*). The remaining lysates (90%) were subjected to co-immunoprecipitation using either an isotype control antibody or anti-Myc antibody to pull down Myc-KIF4. Immunoprecipitates were analyzed by immunoblotting for Myc-KIF4 (*IP; upper panel*) and co-immunoprecipitated HA-NTCP (*IP; lower panel*). Data are representative of three independent experiments.

### A leucine-rich motif (aa 298–301) in the C-terminal region of NTCP is essential for its interaction with KIF4

In addition to NTCP, KIF4 has been reported to mediate the intracellular transport of other host proteins, including human integrin β1 (ITGB1) (25). This raises the possibility that shared functional determinants within these cargo proteins mediate their recognition by KIF4. To identify candidate motifs within the NTCP C-terminal region that may be responsible for KIF4 interaction, we aligned the amino acid sequence of the NTCP C-terminal 116 residues (aa 234–349) with human ITGB1 sequence (Fig. 2A) (Fig. S1). The alignment revealed that the NTCP C-terminal transmembrane domain (C-TMD; aa 281–311) contains a leucine-rich motif (LLLI; aa 298–301), which is also present within the C-TMD of ITGB1 (aa 729–751) at positions 747–750. This conservation suggested that the LLLI motif might function as a determinant for KIF4 recognition. To experimentally evaluate this hypothesis, we generated expression constructs encoding C-terminal HA-tagged NTCP fragments comprising aa 234–349, either wild-type or carrying an alanine-substitution mutation (AAAA) in place of the LLLI motif (Fig. 2B). Each construct was individually co-transfected with a Myc-tagged KIF4 expression plasmid into HEK293FT cells for 72 h, followed by co-IP analysis. Immunoprecipitation with anti-Myc antibody efficiently recovered KIF4, whereas the isotype control yielded no detectable signal, validating the specificity of the assay (Fig. 2C). Both full-length NTCP and the wild-type NTCP (aa 234-349) fragment co-immunoprecipitated with KIF4. In contrast, the AAAA mutant failed to associate with KIF4 (Fig. 2C). These results suggest that the leucine-rich LLLI motif (aa 298–301) within the NTCP C-terminal region is essential for its interaction with KIF4 and likely constitutes a key determinant for KIF4-dependent cargo recognition.

**FIG 2.**
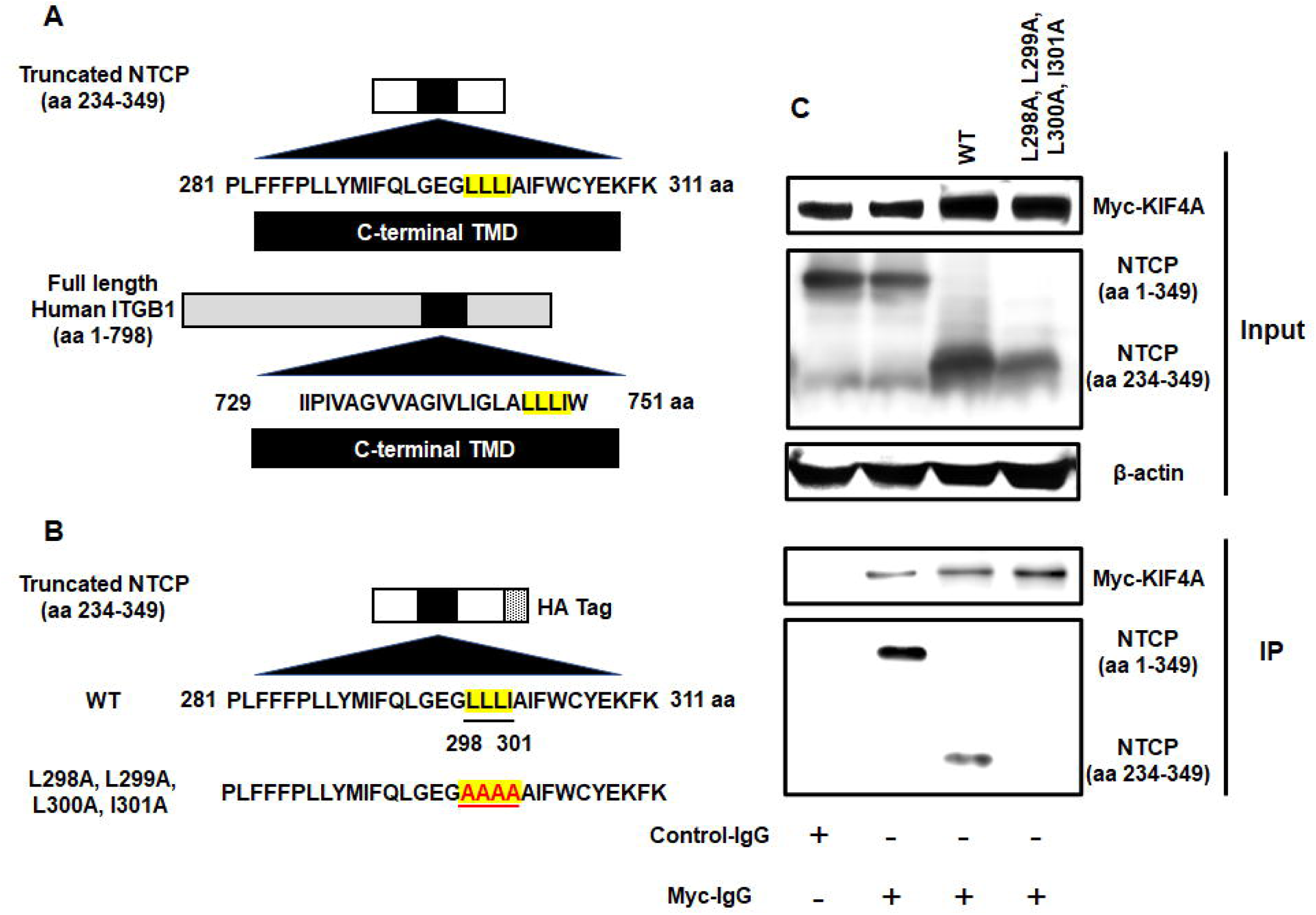
KIF4 recognizes NTCP via a leucine-rich motif (aa 298–301) at the C-terminal end of NTCP. **(A)** Schematic representation of the NTCP truncation mutant comprising the C-terminal 116 residues (aa 234–349) and full-length human ITGB1 (aa 1–798). Black boxes denote the C-terminal transmembrane domains (CT-TMDs) of NTCP (aa 281–311; upper) and ITGB1 (aa 729–751; lower). The shared leucine-rich motif within the CT-TMDs is highlighted in yellow. **(B)** Schematic representation of C-terminally HA-tagged NTCP truncation mutant (aa 234–349) either WT or carrying alanine substitutions at residues L298, L299, L300, and I301. **(C)** HEK293FT cells were co-transfected with expression plasmids encoding Myc-tagged KIF4 and C-terminally HA-tagged full-length NTCP or the NTCP (aa 234–349) truncation mutant, either WT or carrying the indicated alanine substitutions, at a 1:1 ratio. At 72 h post-transfection, cells were lysed, and 10% of each lysate was analyzed by immunoblotting to detect Myc-KIF4 (*Input; uppermost panel*), HA-NTCP (*Input; middle panel*), and β-actin (loading control) (*Input; lowermost panel*). The remaining lysates (90%) were subjected to co-immunoprecipitation using either an isotype control antibody or anti-Myc antibody to pull down Myc-KIF4. Immunoprecipitates were analyzed by immunoblotting for Myc-KIF4 (*IP; upper panel*) and co-immunoprecipitated HA-NTCP (*IP; lower panel*). Data are representative of three independent experiments.

### A conserved leucine-rich motif within the C-terminal region of ITGB1 (aa 747–750) is critical for KIF4-mediated recognition

To further assess the functional importance of the leucine-rich motif (LLLI; aa 747–750) located within the C-TMD of ITGB1 in mediating ITGB1–KIF4 interaction, we constructed expression plasmids encoding C-terminal HA-tagged full-length ITGB1 in either WT form or containing alanine mutations (AAAA) substituting the LLLI motif (Fig. 3A). Each construct was individually co-transfected with a Myc-tagged KIF4 expression vector into HEK293FT cells for 72 h, followed by co-IP analysis. Immunoprecipitation with an anti-Myc antibody efficiently recovered KIF4, whereas no signal was obtained with the isotype control antibody, confirming assay validity (Fig. 3B). Remarkably, WT full-length ITGB1 co-immunoprecipitated with KIF4, whereas the AAAA mutant did not (Fig. 3B). These results indicate that the leucine-rich LLLI motif (aa 747–750) within the ITGB1 C-TMD is crucial for KIF4-dependent recognition and interaction with ITGB1.

**FIG 3.**
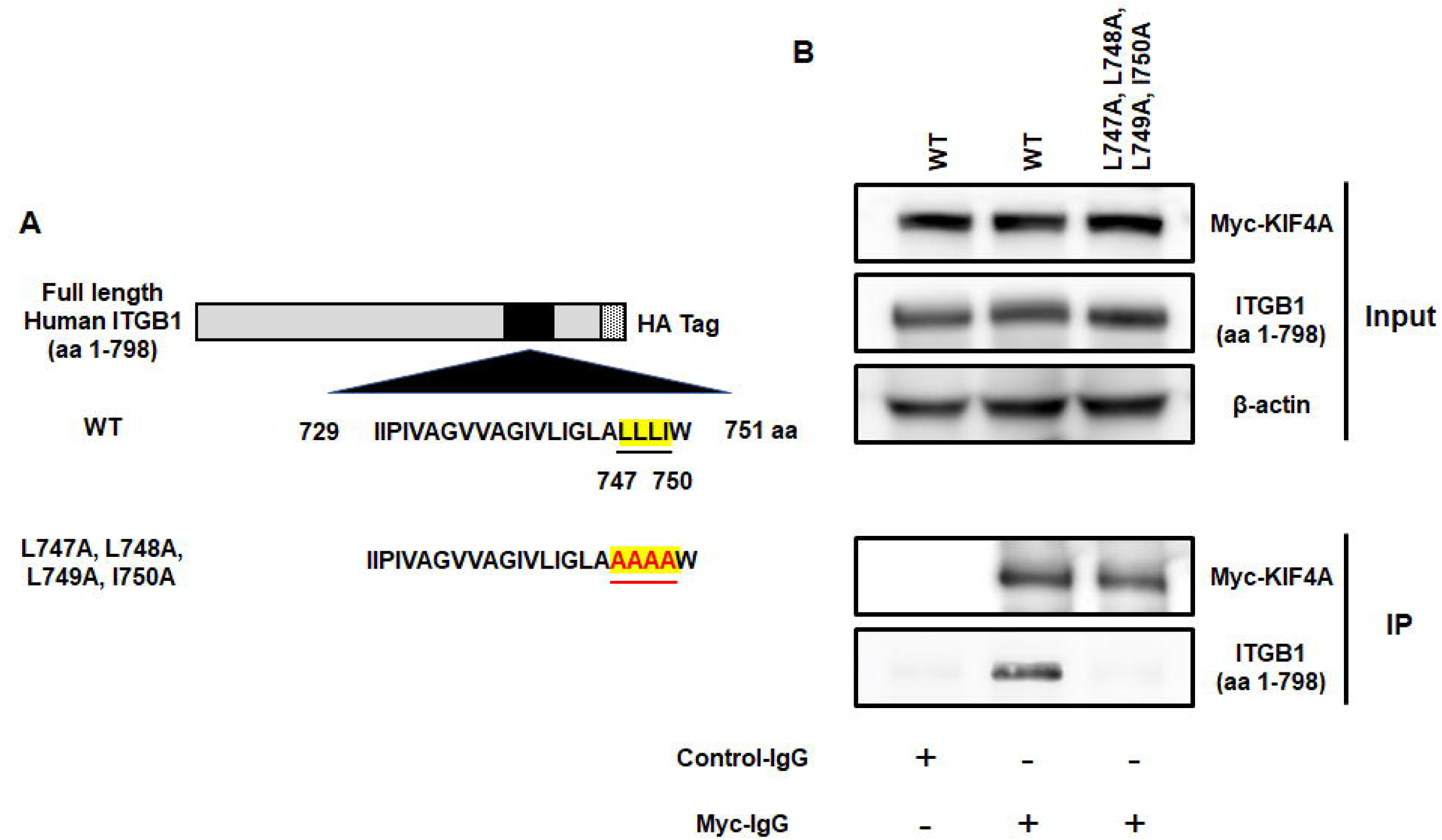
KIF4 selectively recognizes ITGB1 via a leucine-rich motif (amino acids 747–750) at the C-terminal end of ITGB1. **(A)** Schematic representation of C-terminally HA-tagged full-length human ITGB1 (aa 1–798) either WT or harboring alanine substitutions at residues L747, L748, L749, and I750. **(B)** HEK293FT cells were co-transfected with expression plasmids encoding Myc-tagged KIF4 and C-terminally HA-tagged full-length ITGB1 either WT or carrying the indicated alanine substitutions, at a 1:1 ratio. At 72 h post-transfection, cells were lysed, and 10% of each lysate was analyzed by immunoblotting to detect Myc-KIF4 (*Input; uppermost panel*), HA-ITGB1 (*Input; middle panel*), and β-actin (loading control) (*Input; lowermost panel*). The remaining lysates (90%) were subjected to co-immunoprecipitation using either an isotype control antibody or anti-Myc antibody to pull down Myc-KIF4. Immunoprecipitates were analyzed by immunoblotting for Myc-KIF4 (*IP; upper panel*) and co-immunoprecipitated HA-ITGB1 (*IP; lower panel*). Data are representative of three independent experiments.

### The C-terminal leucine-rich motif (aa 298–301) of NTCP is crucial for KIF4-mediated NTCP surface transport and efficient NTCP-dependent HBV/HDV infection

Given that NTCP is transported to the hepatocyte cell surface along microtubules by KIF4 to facilitate HBV and HDV entry (4), we investigated the functional importance of the C-terminal leucine-rich motif of NTCP (aa 298–301) in KIF4-mediated surface NTCP trafficking and HBV/HDV infection. To this end, we created expression plasmids encoding C-terminally HA-tagged full-length NTCP, either as the wild-type (NTCP^WT^) or with alanine substitutions in the LLLI motif (NTCP^AAAA^). HepG2 cells transiently expressing NTCP^WT^ or NTCP^AAAA^ for 72 h were exposed to biotinylation-based surface labeling assay to selectively tag and purify cell surface protein fractions. Although total NTCP expression levels were comparable between the two constructs (Fig. 4A, input), NTCP^AAAA^ exhibited markedly reduced surface localization relative to NTCP^WT^, which predominantly localized to the cell membrane (Fig. 4A, surface). Consistent with impaired surface expression, NTCP^AAAA^ expressing cells displayed significantly reduced binding of the HBV preS1 peptide (*P* **<** 0.001) (Fig. 4B and C). Furthermore, HBV infection was strongly attenuated in NTCP^AAAA^ expressing cells, as evidenced by a significant reduction in intracellular HBV pgRNA levels (*P* **<** 0.001) (Fig. 4D and E) and HBeAg secreted in the cell culture supernatant (*P* **<** 0.001) (Fig. 4F) compared with NTCP^WT^ expressing cells. Similarly, HDV infection was substantially decreased in NTCP^AAAA^ expressing cells, as demonstrated by a marked reduction in HDAg positive cells (*P* **<** 0.001) (Fig. 4G and H). Importantly, the NTCP^AAAA^ substitution did not adversely affect cell viability relative to NTCP^WT^ (Fig. 4I), excluding cytotoxic effects. Collectively, these results demonstrate that the NTCP C-terminal leucine-rich motif (aa 298–301) is essential for KIF4-mediated NTCP surface transport and is consequently critical for efficient HBV and HDV entry.

**FIG 4.**
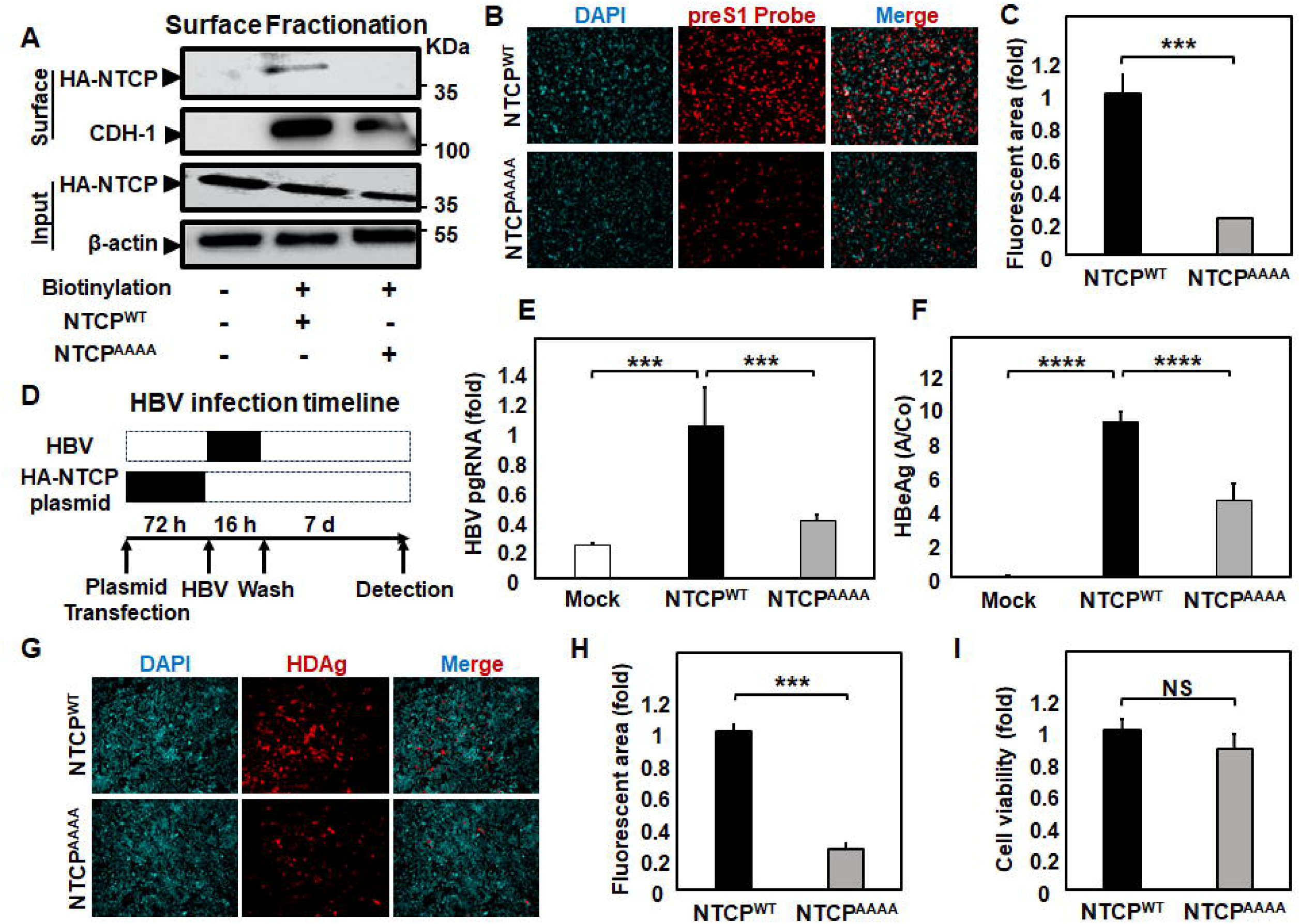
The NTCP C-terminal leucine-rich motif (aa 298–301) is required for KIF4-dependent surface trafficking and HBV/HDV entry. **(A)** HepG2 cells were transfected with plasmids expressing C-terminally HA-tagged full-length NTCP WT (NTCP^WT^) or carrying the L298A, L299A, L300A, I301A substitutions (NTCP^AAAA^). At 72 h post-transfection, the cells were surface-biotinylated or PBS-treated at 4°C for 30 min. prior to cell lysis. Ten percent of each lysate was analyzed by immunoblotting to detect total NTCP (*Input; upper panel*), and β-actin (loading control) (*Input; lower panel*). The remaining lysates were incubated with streptavidin (SA) beads to isolate biotinylated surface proteins. Eluted fractions were analyzed by immunoblotting to detect surface NTCP (*Surface; upper panel*) and CDH-1 (loading control for surface fraction) (*Surface; lower panel*). **(B)** HepG2 cells ectopically expressing NTCP^WT^ or NTCP^AAAA^ were incubated with 40 nM C-terminal TAMRA-labeled and N-terminally myristoylated preS1 peptide (preS1 probe) for 30 min at 37°C; Red and blue signals indicate the preS1 probe binding and nuclei, respectively. **(C)** Quantification of relative fluorescence intensity from panel B. **(D)** Schematic diagram depicting the scheme for plasmid transfection and subsequent HBV infection in HepG2; HepG2 cells were transfected with Mock, NTCP^WT^, or NTCP^AAAA^ plasmids for 72 h, followed by HBV inoculation (10,000 GEq/cell) in the presence of 4% PEG8000 for 16 h. After washing out free viral particles, the cells were cultured for an additional 7 days, followed by detection of different HBV markers. **(E)** Intracellular HBV pgRNA levels were quantified by RT-qPCR and presented as fold changes, relative to NTCP^WT^ transfected cells. **(F)** secreted HBeAg levels in the culture supernatant was measured by ELISA and presented as (A/CO) values. **(G)** HepG2 expressing NTCP^WT^ or NTCP^AAAA^ were inoculated with HDV (100 GEq/cell) in the presence of 5% PEG8000 for 16 h. After washing and removal of the free viral particles, the cells were cultured for 6 days followed by detection of intracellular HDAg by IF; Red and blue signals indicate HDAg and nuclei, respectively. **(H)** Quantification of fluorescence intensity from panel G. **(I)** Cell viability was also assessed by MTT assay. All assays were performed in triplicate, and data from three independent experiments were included. Data are presented as mean ± SD. ***, *P* < 0.001; NS, not significant.

### Structural modeling demonstrates that the NTCP^AAAA^ substitution mutation does not directly alter NTCP’s binding affinity for the preS1 domain

Additionally, we conducted an *in silico* structural analysis using AlphaFold3 (26) to predict the three-dimensional structures of NTCP^WT^, the single alanine-substitution mutants NTCP^L298A^, NTCP^L299A^, NTCP^L300A^, and NTCP^I301A^, and the quadruple mutant NTCP^AAAA^. The overall predicted folds were nearly identical across all variants, with root mean squared deviations (RMSDs) of ≤ 0.3 Å relative to NTCP^WT^ (Fig. 5A, a-f). A close-up view of the mutated C-terminal region likewise revealed no discernible local structural perturbations (Fig. 5A, g). Because cell surface expressed NTCP mediates HBV entry through binding to the preS1 domain of the large hepatitis B surface protein (LHBs) on hepatocytes (3), we next assessed whether the AAAA substitution directly affects NTCP–preS1 interaction. To this end, we employed DDMut-PPI (27), a deep learning–based model that predicts changes in protein–protein interaction binding free energy upon multiple amino acid substitutions (28). The predicted affinity change (ΔΔG^Affinity^ ^WT→L298A, L299A, L300A, I301A^) was -1.11 kcal/mol (root mean squared error: 1.33 kcal/mol), predicting no substantial reduction in the intrinsic binding affinity between NTCP and the HBV preS1 domain (Fig. 5B and C). Collectively, these *in silico* analyses support the conclusion that the impaired ability of the NTCP^AAAA^ mutant to support HBV and HDV infection results from defective surface trafficking rather than from a direct disruption of NTCP–preS1 binding.

**FIG 5.**
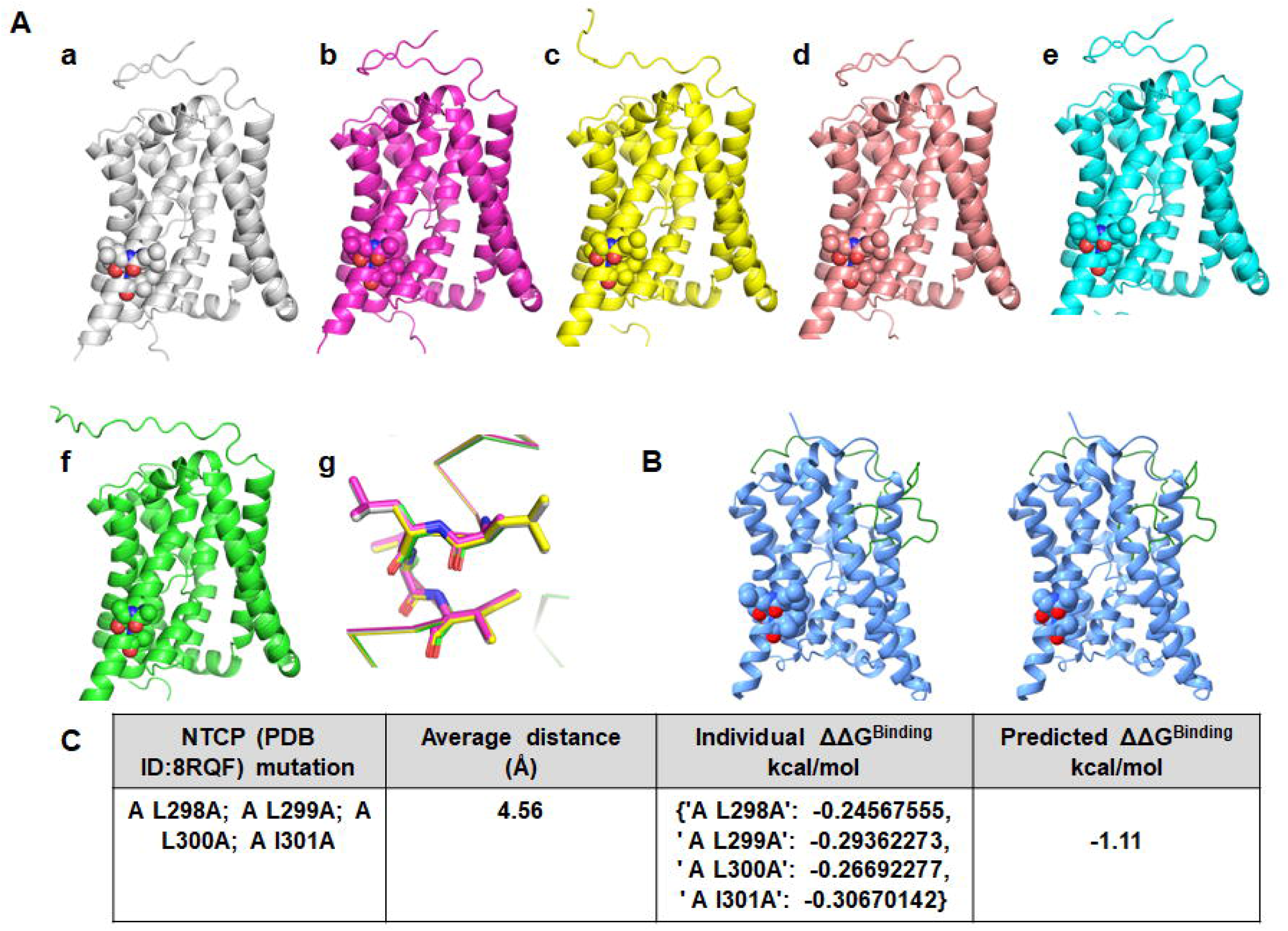
*In silico* structural modeling suggests the NTCP^AAAA^ substitution mutation does not alter NTCP/preS1 binding. **(A)** Panels A (a–f) show the predicted human NTCP structures of the wild type and the single alanine mutants; L298A, L299A, L300A, and I301A, and the quadruple mutant L298A/L299A/L300A/I301A, respectively. Panel A (g) overlays the structures at the mutation sites. The colors for L298A, L299A, L300A, I301A, and L298A/L299A/L300A/I301A are magenta, yellow, tint, cyan, and green, respectively. (B) Panel B (a, b) shows the predicted human NTCP and PreS1 structures of the wild type (a) and the quadruple mutant L298A/L299A/L300A/I301A (b), respectively. **(C)** Predicted effects of single and quadruple alanine substitutions within the NTCP leucine-rich motif on binding free energy between NTCP and the HBV preS1 domain. Individual ΔΔG^Binding^ values (kcal/mol) are shown for the single mutants (L298A, L299A, L300A, I301A), along with the predicted ΔΔG^Binding^ (kcal/mol) for the quadruple mutant (L298A/L299A/L300A/I301A).

### L298 and L299 of NTCP constitute the minimal critical residues required for mediating the interaction between NTCP and KIF4

As our results indicated that the NTCP leucine-rich motif (aa 298–301) is required for its interaction with KIF4 and subsequent surface transport, we next sought to define the minimal residues within this motif that mediate the interaction. To this end, we generated expression plasmids encoding C-terminal HA-tagged NTCP fragments spanning aa 234–349, either wild-type or carrying a single alanine substitution at L298, L299, L300, or I301 (Fig. 6A). Each construct was co-transfected with a Myc-tagged KIF4 expression plasmid into HEK293FT cells for 72 h, followed by co-IP analysis. Immunoprecipitation with an anti-Myc antibody efficiently recovered KIF4, whereas the isotype control yielded no detectable signal, validating the assay specificity (Fig. 6B). The wild-type NTCP fragment (aa 234–349), as well as the L300A and I301A variants, robustly co-immunoprecipitated with KIF4. In contrast, fragments carrying L298A or L299A exhibited absent or markedly reduced co-IP signals. As expected, the AAAA mutant fragment used as a negative control failed to associate with KIF4 (Fig. 6B), further validating the co-IP specificity. Taken together, these findings indicate that NTCP residues L298 and L299 represent the minimal and essential determinants required for efficient NTCP–KIF4 interaction.

**FIG 6.**
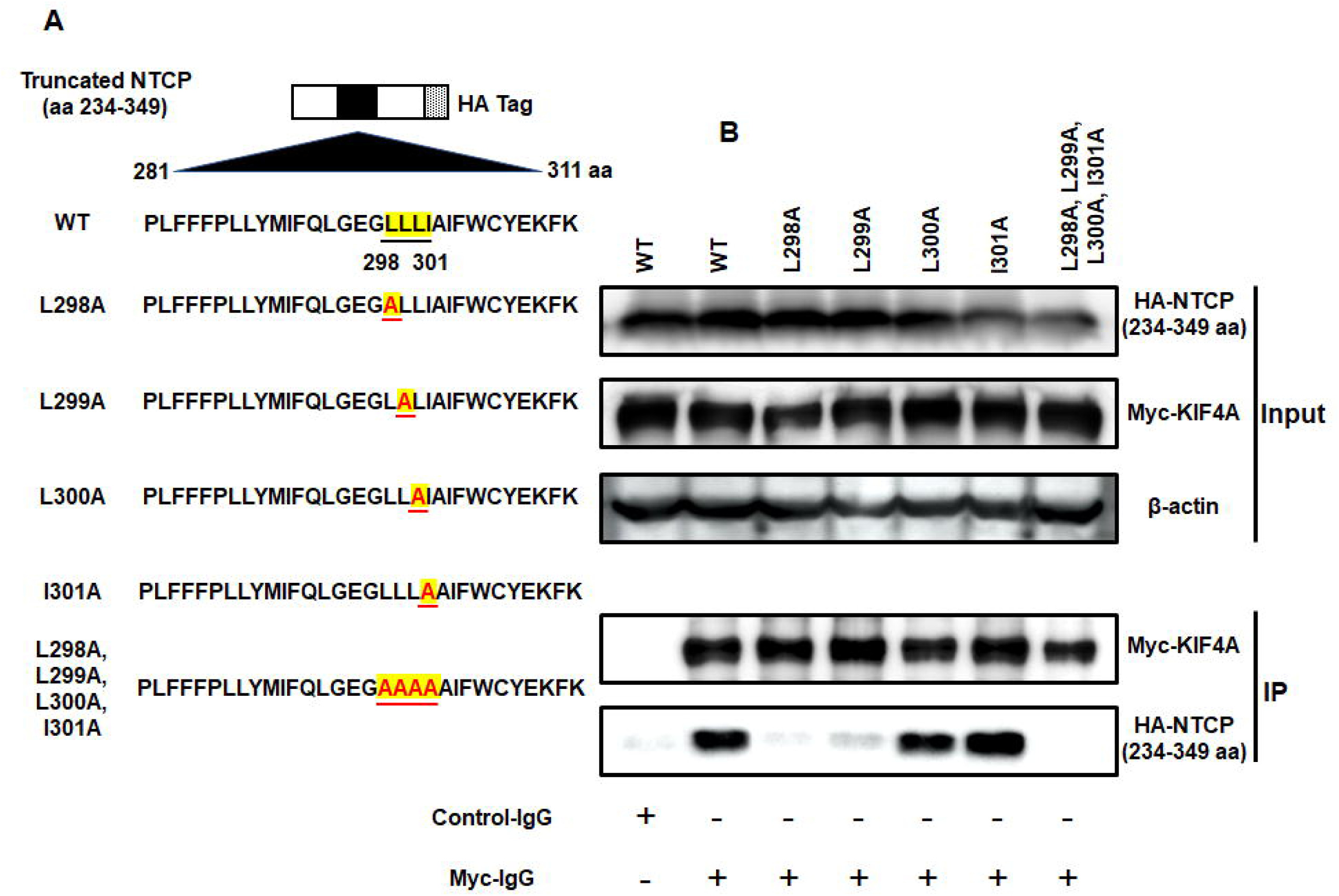
NTCP residues L298 and L299 are the minimal determinants essential for interaction with KIF4. **(A)** Schematic representation of C-terminally HA-tagged NTCP truncation mutant (aa 234–349) either WT, carrying single alanine substitutions at residues L298, L299, L300, or I301, or quadruple alanine substitutions (L298A/L299A/L300A/I301A). **(B)** HEK293FT cells were co-transfected with expression plasmids encoding Myc-tagged KIF4 and C-terminally HA-tagged NTCP (aa 234–349) truncation mutant, either WT or carrying the indicated alanine substitutions, at a 1:1 ratio. At 72 h post-transfection, cells were lysed, and 10% of each lysate was analyzed by immunoblotting to detect HA-NTCP variants (*Input; uppermost panel*), Myc-KIF4 (*Input; middle panel*), and β-actin (loading control) (*Input; lowermost panel*). The remaining lysates (90%) were subjected to co-immunoprecipitation using either an isotype control antibody or anti-Myc antibody to pull down Myc-KIF4. Immunoprecipitates were analyzed by immunoblotting for Myc-KIF4 (*IP; upper panel*) and co-immunoprecipitated HA-NTCP (*IP; lower panel*). Data are representative of three independent experiments.

## Discussion

ATP-dependent intracellular transport of cargo proteins along microtubule networks by motor proteins is a fundamental process required for cellular homeostasis. Microtubule-based motors, primarily kinesins and dyneins, are ATP-driven transporters that mediate the directional trafficking of diverse intracellular cargos. Kinesins generally drive anterograde transport toward the cell periphery, whereas dyneins and certain C-terminal kinesins facilitate retrograde movement toward the perinuclear region (10). A growing body of evidence demonstrates that these motor proteins play critical roles in viral infection by directly interacting with viral components, thereby promoting viral replication and propagation (29–32). In addition, we and others have shown that viruses can indirectly exploit the kinesin transport machinery by engaging host proteins that are transported by motor proteins and subsequently co-opted to support viral entry or replication (4, 25, 33).

Notably, the motor protein KIF4 interacts with and anterogradely transports several host proteins, including human sodium taurocholate co-transporting polypeptide (NTCP), to the hepatocyte cell surface, where NTCP serves as an entry receptor for HBV and HDV. KIF4 has also been reported to transport integrin β1 (ITGB1) and L1 cell adhesion molecule (L1CAM)–containing vesicles in developing cortical neurons (4, 7, 25). These observations raise the question of how KIF4 selectively recognizes and transports specific host cargos. Structurally, KIF4 contains an N-terminal ATPase motor domain responsible for microtubule engagement and a C-terminal cargo-binding tail that mediates cargo recognition (10). In the present study, we used NTCP and ITGB1 as model cargos to interrogate the molecular basis of KIF4-mediated transport.

Through a combination of amino acid sequence alignment and biochemical analyses, we identified a conserved C-terminal leucine-rich motif in both NTCP (aa 298–301) and ITGB1 (aa 747–750) that is essential for cargo recognition and transport mediated by KIF4. NTCP is a liver-specific, basolaterally expressed transporter that mediates the hepatic uptake of bile acids and functions as the bona fide entry receptor for HBV and HDV viruses (3, 34). In this study, alanine substitution of the NTCP leucine-rich motif (aa 298–301) disrupted normal NTCP surface localization, markedly reduced binding of a fluorescent HBV preS1 probe, and significantly attenuated HBV and HDV infection (Fig. 4). These findings underscore the functional importance of this motif in KIF4-mediated NTCP trafficking to the cell surface.

Although several NTCP orthologs from different species support HBV binding and infection (35, 36), sequence alignment of the C-terminal transmembrane sequence (aa 294–305) revealed that the putative leucine-rich motif (aa 298–301) is not strictly conserved among HBV-susceptible species (Fig. S3). The fact that this motif is not fully conserved among different species supporting HBV infection suggests it does not play a direct role in mediating NTCP binding to the HBV preS1 domain. Instead, it may possess a function unique to the host cell, most likely in regulating the intracellular trafficking of NTCP, rather than directly facilitating viral binding to NTCP. Consistent with this interpretation, *in silico* structural modeling using AlphaFold supported that alanine substitution of this motif does not induce detectable global or local conformational changes in NTCP (Fig. 5). Comparison with the experimentally determined human NTCP cryo-EM structure (PDB: 7PQG) showed that all AlphaFold3 models most closely resembled the inward-facing conformation, with an RMSD of approximately 1.5 Å (34), and no mutation-associated structural perturbations were predicted. This apparent structural insensitivity may reflect inherent limitations of machine-learning–based prediction, which tends to favor conformations represented in the training data, or alternatively suggests that NTCP may adopt distinct, context-dependent conformational states influenced by membrane environment or interacting partners that are not captured by isolated *in silico* modeling. In parallel, DDMut-PPI analysis predicted only a minimal change in binding free energy between NTCP and the HBV preS1 domain upon introduction of the AAAA mutation, further supporting the conclusion that this motif does not directly contribute to intrinsic NTCP–preS1 binding affinity. While these computational predictions cannot substitute for direct biochemical binding measurements, they are consistent with prior studies identifying alternative NTCP residues as the primary determinants of HBV preS1 engagement and viral internalization (37–41).

Taken together with the comparable steady-state expression levels of NTCP WT and the AAAA mutant (Fig. S2) and the markedly reduced surface localization of the AAAA variant, indicative of impaired motor-dependent transport, these data support a trafficking defect—rather than an intrinsic loss of receptor function—as the primary basis for the reduced NTCP-mediated HBV and HDV infectivity observed for the AAAA mutant. Collectively, our combined biochemical, structural, and computational analyses support a model in which the NTCP leucine-rich motif is dispensable for direct viral binding but is essential for KIF4-dependent NTCP recognition and surface transport, thereby indirectly regulating HBV and HDV entry.

Integrin β1 (ITGB1/CD29) is a transmembrane protein that forms functional heterodimers with multiple α-integrin subunits, including ITGA2 and ITGA3, thereby generating receptors that are essential for cellular homeostasis. Through binding to extracellular matrix (ECM) components such as laminin, collagen, and fibronectin, ITGB1-containing integrins physically link the ECM to the intracellular cytoskeleton and regulate key cellular processes, including adhesion, proliferation, migration, and gene expression (42, 43). Beyond these physiological functions, ITGB1 heterodimers also serve as entry factors for a variety of viruses (43–47) and contribute to tumor invasion and metastasis by enabling cancer cells to adhere to the basement membrane of target tissues (48). Collectively, these properties position ITGB1 as an attractive antiviral and anticancer target. In this context, elucidating the molecular mechanisms determining KIF4-dependent ITGB1 transport may provide a strategy to selectively disrupt ITGB1 surface delivery without broadly impairing other KIF4-dependent cellular functions, thereby offering a potential avenue for the development of targeted antiviral and anticancer therapies.

Here, we provide the first mechanistic insight into how the motor protein KIF4 selectively recognizes specific host cargo proteins for directed trafficking to the cell surface. Our findings reveal that surface protein delivery is a highly organized and tightly regulated process, in which defined sequence determinants enable selective engagement with distinct motor proteins to ensure precise spatial localization. This work advances our understanding of motor-driven secretory pathways and highlights cargo-specific recognition as a key principle governing intracellular transport to the plasma membrane.

## Acknowledgement

We gratefully acknowledge Dr. John Taylor at the Fox Chase Cancer Center, the USA for providing pSVLD3 plasmid, Toru Hirota at JFCR, Japan for providing Myc-tagged KIF4; Dr. Hiroyuki Miyoshi at RIKEN, Japan, for providing HA-tagged NTCP. This study was supported by Grants in aid for Scientific Research C, 23K06575 and the Research Program on Hepatitis B from AMED (grants number: 26fk0310532s0102; and 26fk0310528s0102) to Hussein H. Aly, and the Research Program on Hepatitis B from AMED (grant number: 25fk0310532h0001) to Kazuaki Chayama.

## Supplementary figure legends

**FIG S1**

**Sequence alignment of the NTCP C-terminal region with human ITGB1.** Alignment of the C-terminal 116 amino acids of NTCP (aa 234–349) with human ITGB1. Conserved residues and regions of sequence similarity are marked or highlighted.

**FIG S2**

**Protein expression of NTCP and ITGB1 (WT *vs* quadruple alanine mutant) in HEK293 FT cells.** HEK293FT cells were transfected with mock plasmid or plasmids expressing C-terminally HA-tagged full-length NTCP WT, NTCP carrying L298A/L299A/L300A/I301A substitutions, C-terminally HA-tagged full-length ITGB1 WT, or ITGB1 harboring L747A/L748A/L749A/I750A substitutions. At 72 h post-transfection, cells were lysed and analyzed by immunoblotting to detect NTCP or ITGB1 (*upper panel*) and β-actin (*lower panel*; loading control). Data are representative of three independent experiments. Data are representative of three independent experiments.

**FIG S3**

**Sequence alignment of NTCP residues 294–305 within the C-terminal transmembrane region across HBV-susceptible species.** The C-terminal transmembrane segment (aa 294–305) of NTCP orthologs from human (*Homo sapiens*), cat (*Felis catus*), rabbit (*Oryctolagus cuniculus*), ferret (*Mustela putorius furo*), woodchuck (*Marmota marmota*), big brown bat (*Eptesicus fuscus*), aardvark (*Orycteropus afer afer*), horse (*Equus caballus*), chinese tree shrew (*Tupaia chinensis*), whale (*Balaenoptera acutorostrata scammony*) was shown. Residues corresponding to the putative leucine-rich motif identified in human NTCP are highlighted in purple.

