## Supplementary material for "Identification of molecular determinants governing KIF4 cargo recognition of NTCP and their role in HBV/HDV infection": Fig. S1

### Sup. 1

```
                                       1
Human NTCP (234-349 aa)    ......FLLGYV..........................................................
Human ITGB1                MNLQPIFWIGLISSVCCVFAQTDENRCLKANAKSCGECIQAGPNCGWCTNSTFLQEGMPTSARCDDLEAL

Human NTCP (234-349 aa)    ......................................................................
Human ITGB1                KKKGCPPDDIENPRGSKDIKKNKNVTNRSKGTAEKLKPEDITQIQPQQLVLRLRSGEPQTFTLKFKRAED

Human NTCP (234-349 aa)    ......................................................................
Human ITGB1                YPIDLYYLMDLSYSMKDDLENVKSLGTDLMNEMRRITSDFRIGFGSFVEKTVMPYISTTPAKLRNPCTSE

Human NTCP (234-349 aa)    ...................................................................LSA
Human ITGB1 (1-798 aa)     QNCTSPFSYKNVLSLTNKGEVFNELVGKQRISGNLDSPEGGFDAIMQVAVCGSLIGWRNVTRLLVFSTDA

                           10                  20
Human NTCP (234-349 aa)    LF.............CLNGRCRRTVSME..........................................
Human ITGB1 (1-798 aa)     GFHFAGDGKLGGIVLPNDGQCHLENNMYTMSHYYDYPSIAHLVQKLSENNIQTIFAVTEEFQPVYKELKN

                                                                                          30
Human NTCP (234-349 aa)    ..............................................................TGCQNVQL
Human ITGB1 (1-798 aa)     LIPKSAVGTLSANSSNVIQLIIDAYNSLSSEVILENGKLSEGVTISYKSYCKNGVNGTGENGRKCSNISI

Human NTCP (234-349 aa)    CS......TI............................................................
Human ITGB1 (1-798 aa)     GDEVQFEISITSNKCPKKDSDSFKIRPLGFTEEVEVILQYICECECQSEGIPESPKCHEGNGTFECGACR

Human NTCP (234-349 aa)    ......................................................................
Human ITGB1 (1-798 aa)     CNEGRVGRHCECSTDEVNSEDMDAYCRKENSSEICSNNGECVCGQCVCRKRDNTNEIYSGKFCECDNFNC

Human NTCP (234-349 aa)    ......................................................................
Human-ITGB1 (1-798 aa)     DRSNGLICGGNGVCKCRVCECNPNYTGSACDCSLDTSTCEASNGQICNGRGICECGVCKCTDPKFQGQTC

Human NTCP (234-349 aa)    ......................................................................
Human ITGB1 (1-798 aa)     EMCQTCLGVCAEHKECVQCRAFNKGEKKDTCTQECSYFNITKVESRDKLPQPVQPDPVSHCKEKDVDDCW

                                          40        50            60               70        80
Human NTCP (234-349 aa)    ..............LNVA.FPPEVIGPLFFFPL....LYMIFQLGEGLLLI.......AIFWCYEKFKTP
Human ITGB1 (1-798 aa)     FYFTYSVNGNNEVMVHVVENPECPTGPDI.IPIVAGVVAGIVLIGLALLLIWKLLMIIHDRREFAKFEKE

                                        90        100        110
Human NTCP (234-349 aa)    KDK......TKMIYTAATTEETIPGALGNGTYKGEDCSPCTA
Human ITGB1 (1-798 aa)     KMNAKWDTGENPIYKSAVTT......VVNPKYEGK.......
```
