## Supplementary figures and images for "Identification of molecular determinants governing KIF4 cargo recognition of NTCP and their role in HBV/HDV infection"

### Fig. S2

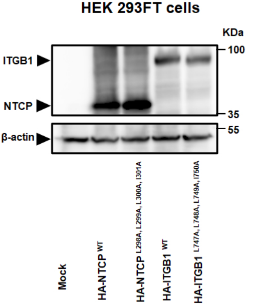

### Fig. S3

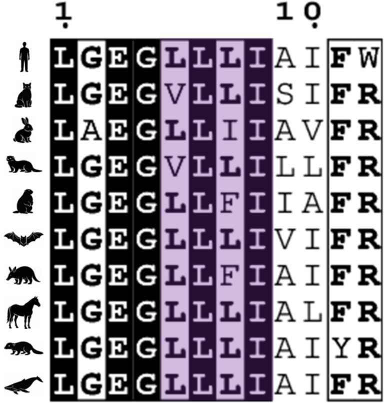
